# Serum metabolomic analysis of men on a low-carbohydrate diet for biochemically recurrent prostate cancer reveal the potential role of ketogenesis to slow tumor growth: A secondary analysis of the CAPS2 diet trial

**DOI:** 10.1101/2021.12.29.474437

**Authors:** Jen-Tsan Chi, Pao-Hwa Lin, Vladimir Tolstikov, Lauren Howard, Emily Y. Chen, Valerie Bussberg, Bennett Greenwood, Niven R. Narain, Michael A. Kiebish, Stephen J. Freedland

## Abstract

**Background:** Systemic treatments for prostate cancer (PC) have significant side effects. Thus, newer alternatives with fewer side effects are urgently needed. Animal and human studies suggest the therapeutic potential of low carbohydrate diet (LCD) for PC. To test this possibility, Carbohydrate and Prostate Study 2 (CAPS2) trial was conducted in PC patients with biochemical recurrence (BCR) after local treatment to determine the effect of a 6-month LCD intervention vs. usual care control on PC growth as measured by PSA doubling time (PSADT). We previously reported the LCD intervention led to significant weight loss, higher HDL, and lower triglycerides and HbA1c with a suggested longer PSADT. However, the metabolic basis of these effects are unknown.

**Methods:** To identify the potential metabolic basis of effects of LCD on PSADT, serum metabolomic analysis was performed using baseline, month 3, and month 6 banked sera to identify the metabolites significantly altered by LCD and that correlated with varying PSADT.

**Results:** LCD increased the serum levels of ketone bodies, glycine and hydroxyisocaproic acid. Reciprocally, LCD reduced the serum levels of alanine, cytidine, asymmetric dimethylarginine (ADMA) and 2-oxobutanoate. As high ADMA level is shown to inhibit nitric oxide (NO) signaling and contribute to various cardiovascular diseases, the ADMA repression under LCD may contribute to the LCD-associated health benefit. Regression analysis of the PSADT revealed a correlation between longer PSADT with higher level of 2-hydroxybutyric acids, ketone bodies, citrate and malate. Longer PSADT was also associated with LCD reduced nicotinamide, fructose-1, 6-biphosphate (FBP) and 2-oxobutanoate.

**Conclusion:** These results suggest a potential association of ketogenesis and TCA metabolites with slower PC growth and conversely glycolysis with faster PC growth. The link of high ketone bodies with longer PSADT supports future studies of ketogenic diets to slow PC growth.

## Introduction

Prostate cancer (PC) is the most common non-cutaneous cancer and second leading cause of cancer death in the U.S (1). For men with non-metastatic PC who require treatment, local therapy is not always curative and they may progress to needing systemic therapies, such as androgen-deprivation therapy (ADT), chemotherapy and androgen-targeted therapy (2, 3). However, these systemic therapies are often associated with significant side effects. Furthermore, our team and others found obese men are more likely to present with high-grade PC, progress to metastases, and die from PC (4–9). Thus, newer alternatives with fewer side effects that compliment standard of care are urgently needed, especially in overweight and obese men, which is more than 2/3rds of American men (10).

In two PC xenografts comparing western diet and a low carbohydrate diet (LCD), we found LCD, even without weight loss, prolonged survival (10, 11) and slowed PC growth in a transgenic mouse model (12). In the Carbohydrate and Prostate Study 2 (CAPS2) trial (13), we tested the effect of an 6-month LCD intervention vs. usual care control on PC growth in men with biochemical recurrence after definitive therapy for PC. PC growth was measured by prostate specific antigen (PSA) doubling time (PSADT), a measure that is strongly correlated with death in men with recurrent PC(14). We found that the LCD intervention resulted in significant weight loss, improvements in HDL, and reductions in triglycerides and HbA1c with a suggested longer PSADT. Even though future larger studies are needed to verify the benefits of a LCD with more reliable surrogates of tumor growth and cancer-specific mortality, such as metastasis-free survival (13), this study also provided the opportunity to explore potential metabolic mechanisms linking LCD and PC progression.

One powerful approach to understand the metabolic effects of LCD is to profile all the serum small molecule metabolites by nuclear magnetic resonance (NMR) or mass spectrometry (MS). Such metabolomic approaches can serve as an integrated read-out of all the upstream somatic mutations, gene expression protein and biochemical activities resulting from the effects of dietary intervention. For example, the metabolomic profiling of The Cancer Genome Atlas breast dataset can be interpreted in the context of various intrinsic subtypes and somatic mutations (15, 16). The metabolomic analysis of PC also reveals dysregulated sarcosine-related pathways (17, 18). Recently, we reported the serum metabolomic analysis of PC patients in the control arm of the CAPS1 trial after 3 and 6 months of ADT (19). Such analysis revealed that ADT reduced androsterone sulfate, 3-hydroxybutyric acid, acyl-carnitines while increased serum glucose. Since ADT reduced ketogenesis, metabolomic analysis also showed that LCDs reversed many ADT-induced metabolic abnormalities including reducing ketogenesis without affecting the ADT-reduced androsterone sulfate (20). Several studies have employed metabolomic approaches to analyze the effects of LCD (21, 22). However, the metabolomic effects of LCD in PC patients who are not receiving concomitant systemic treatment remains unknown. In the current study, we performed serum metabolomic analysis of men enrolled in the CAPS2 trial to determine the effects of LCD on serum metabolome and how the serum metabolomic responses relate to PSADT to identify metabolic links between LCD and slowed PC growth.

## Participants and Methods

### Study Design

The primary results of the CAPS2 have previously been published (13). In brief, The CAPS2 was a multicenter phase II randomized controlled trial (RCT) of LCD vs. control diet (asked to make no diet changes). Key eligibility included biochemical recurrence after local PC treatment, BMI≥24 kg/m2, PSA within the past two months between 3 and 20 ng/ml (prior radiation) or between 0.4 and 20 ng/ml (prior prostatectomy), PSADT 3-36 months, no metastases, not using weight loss treatment and not already following an LCD. Each patient signed a written consent. After confirming eligibility, patients completed a baseline visit and were randomized 1:1 to receive the LCD intervention or a no-dietary intervention control for 6 months. LCD patients were coached by a dietitian by phone for six months to restrict carbohydrates to ≤20grams/day. Controls were instructed to avoid dietary changes. The main outcome was PSADT assed at the end of intervention. Detailed description of the trial has been published previously (13).

### Data Collection and analysis

Data collection occurred at baseline (BL), 3-(M3), and 6-month (M6) post randomization. At each visit, weight (without shoes and in light clothing) was measured, and fasting blood was collected for PSA measurement I commercial laboratories (LabCorp for Duke and Durham Veteran Affairs sites and Cedars-Sinai clinical lab for the Cedars-Sinai site). PSA values were used to calculate PSADT as the natural log of 2 (0.693) divided by the log slop of the linear regression line of log of the PSA over time(14). Fasting blood samples were used for metabolomics analysis utilizing GC/MS-TOF (Gas chromatography–Mass Spectrometry Time of Flight analyzer), QqQ LC (Liqu Chromatography)-HILIC (Hydrophilic interaction chromatography)-MS/MS, and TripleTOF LC-RP-MS as described previously (19, 23).

### Metabolomics Analysis

Serum samples for metabolomics analysis were performed by Berg (Berg LLC, Framingham, MA) as previously described (19, 24–27). In brief, the metabolites were extracted using a 3:3:2 v/v mixture of acetonitrile, isopropanol, and water. Extracts were divided into three parts: 75 uL for gas chromatography combined with time-of-flight high-resolution MS, 150 uL for reversed-phase LC-MS, and 150 uL for hydrophilic interaction chromatography with LC/MS-MS as previously described (19, 24–27). NEXERA XR UPLC system (Shimadzu, Columbia, MD, USA) was used with the Triple Quad 5500 System (AB Sciex, Framingham, MA, USA) to perform hydrophilic interaction LC analysis. NEXERA XR UPLC system (Shimadzu, Columbia, MD, USA), coupled with the Triple TOF 6500 System (AB Sciex, Framingham, MA, USA) to perform reversed-phase LC analysis, and Agilent 7890B gas chromatograph (Agilent, Palo Alto, CA, USA) interfaced to a TOF Pegasus HT MS (Leco, St. Joseph, MI, USA). The GC system was fitted with a Gerstel temperature-programmed injector, cooled injection system (model CIS 4). An automated liner exchange (ALEX) (Gerstel, Muhlheim an der Ruhr, Germany) was used to eliminate cross-contamination between sample runs. Quality control was performed using metabolite standards mixture and pooled samples as preciously described (24, 27). A quality control sample containing a standard mixture of amino and organic acids (Sigma-Aldrich) as described (25, 26). A pooled quality control sample was obtained by taking an aliquot of all samples from the study to determine the optimal dilution of the batch samples, validate metabolite identification and peak integration. Collected raw data was manually inspected, merged, imputed and normalized to calculate the changes of metabolites in all the specimen. Metabolite identification was performed using in house authentic standards analysis. Metabolite annotation was used utilizing recorded retention time and retention indexes, recorded MSn and HRAMSn data matching with METLIN, NIST MS, Wiley Registry of Mass Spectral Data, HMDB, MassBank of North America, MassBank Europe, Golm Metabolome Database, SCIEX Accurate Mass Metabolite Spectral Library, MzCloud, and IDEOM databases.

### Metabolite pathway analysis

Metabolomic data were analyzed as previously described (26). Identified metabolites were subjected to pathway analysis with MetaboAnalyst 4.0, using Metabolite Set Enrichment Analysis (MSEA) module which consists of an enrichment analysis relying on measured levels of metabolites and pathway topology and provides visualization of the identified metabolic pathways. Accession numbers of detected metabolites (HMDB, PubChem, and KEGG Identifiers) were generated, manually inspected, and utilized to map the canonical pathways. MSEA was used to interrogate functional relation, which describes the correlation between compound concentration profiles and clinical outcomes.

### Statistical Analysis

Significant changes were examined by ANOVA and shown in heatmaps and box plots. Pearson correlation was conducted to examine associations between PSADT and metabolites. Volcano plots were used to visually examine the changes in metabolites at M3 and M6 from baseline. The changes of metabolites at M3 and M6 were normalized against the BL using Python scripts (supplemental file 1) modified from previous studies of transcriptional response (28–31) to derive the LCD-induced changes of all metabolites at M3 and M6 from corresponding BL in the control and LCD arms. Given the small number of men included in the study, no formal power calculations were performed for this report and all results are considered hypothesis-generating.

## Results

### Serum metabolites altered after 3 and 6 months of LCD diets

In this secondary analysis to the CAPS2 study, we measured and identified serum metabolites that were altered by LCD at M3 and M6 and correlated with the variations in the PSADT (Fig 1). Specifically, we identified serum metabolites which have significant changes between BL (immediately before dietary changes) and after 3 (M3) and 6 months (M6) of LCD diets. Of the 45 men who completed CAPS2, 39 had metabolite data available at baseline, 36 at M3, and 33 at M6. Among the 489 metabolites identified in the serum samples, 27 metabolites were identified showing at least a 1.2-fold change and t-tests p<0.05 at M3 in the LCD arm (Fig 2A, Supplemental Table 1). The top metabolites, as shown in volcano plots (Fig 2A, Supplemental Table 1), included an increase in 3-hydroxybutyric acid, 3-hydroxy-2-methylbutyriuc acid, 2-hydroxybutyric acid and 2-hydroxyvalerate. These changes in ketone bodies are the expected metabolic effects of LCD and indicate the dietary adherence of the participants. In addition, there was an increase in glycine and isobutyl-glycine and reduced levels of alanine, asymmetric dimethylarginine (ADMA), proline betaine and 2-phenylglycine (Fig 2A, Supplemental Table 1). Using similar filtering criteria (1.2-fold change and t-tests p<0.05), 53 metabolites were identified as significantly changed between BL and M6 in LCD as shown in volcano plots (Fig 2B, Supplemental Table 2). At M6, the top changes included an increase in glycine, 3-hydroxybutyric acid, 3-hydroxy-2-methylbutyriuc acid and 2-hydroxyvalerate. There was also a reduction in alanine, 2-oxobutanoate, proline betaine and ADMA (Fig 2B, Supplemental Table 2).

**Figure 1:**
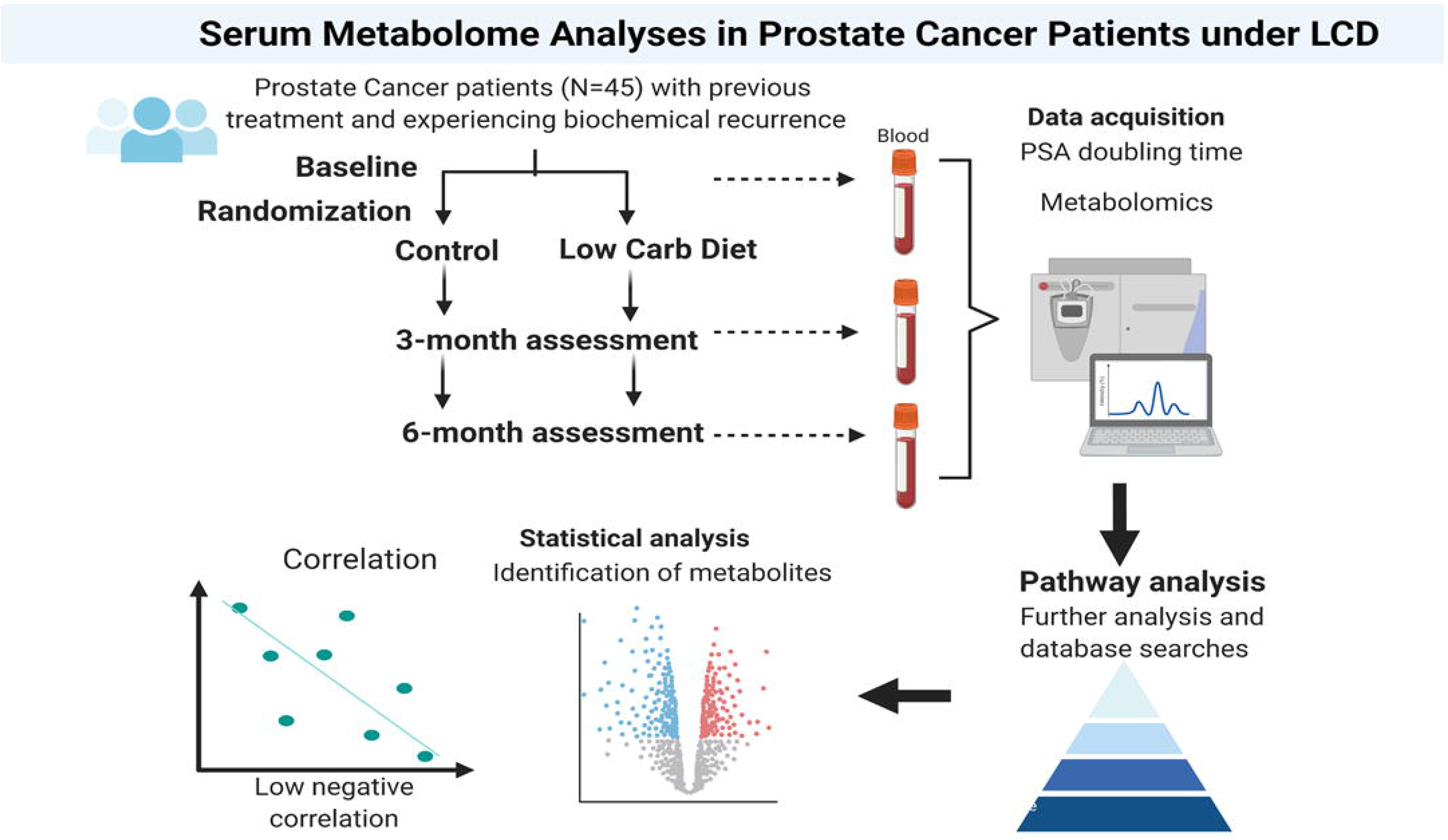
Overview of the experimental design of the study.

**Figure 2:**
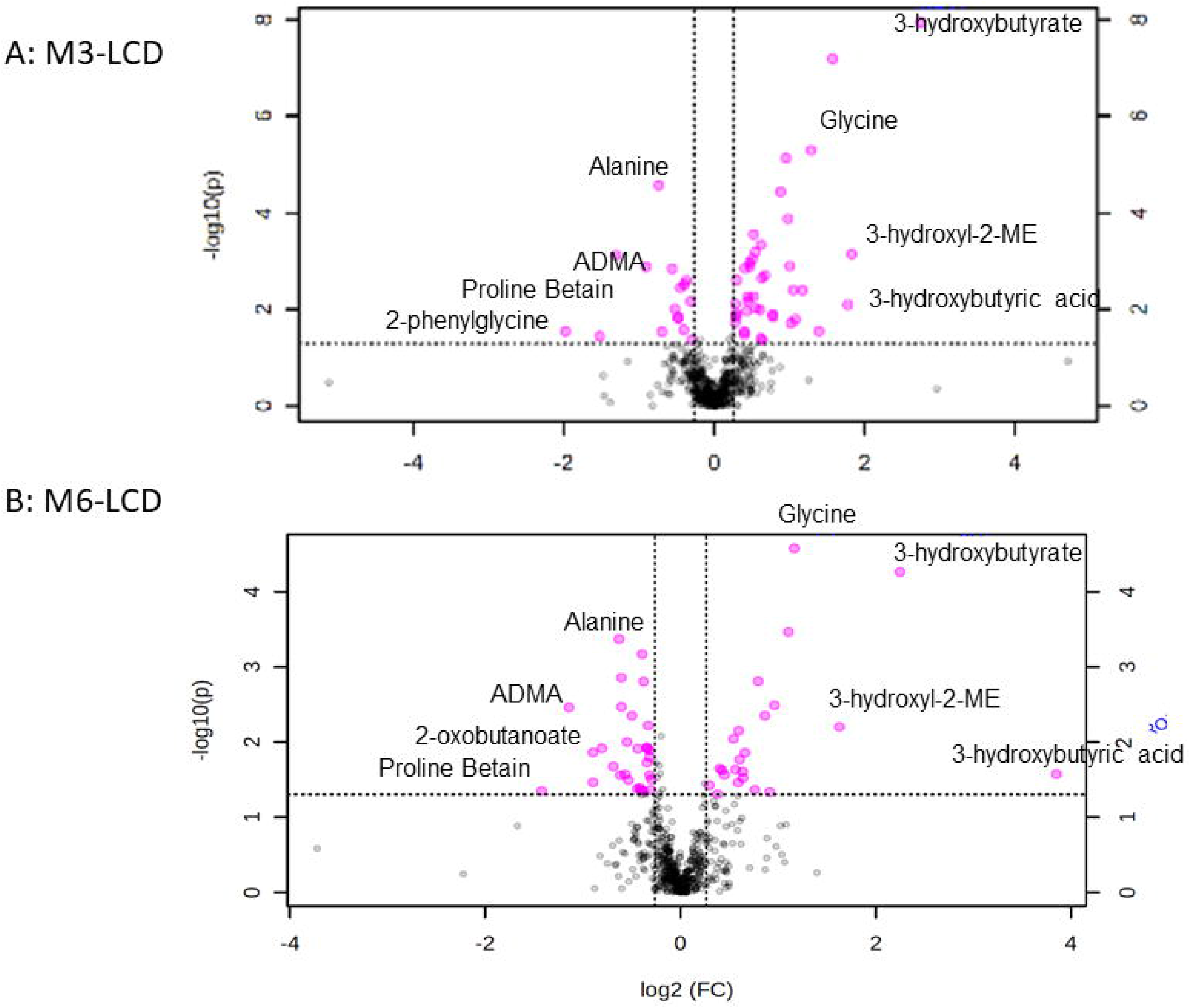
Top serum metabolites altered at 3- and 6-month in participants receiving LCD. (A) Metabolites were selected by Volcano plot analysis of pairwise comparison between the BL vs. M3 (A) and BL vs. M6 (B) of LCD. Selected metabolites were indicated.

### Serum metabolic pathways affected by LCD at 3 and 6 months

As several LCD-affected metabolites belong to similar metabolic pathways, we employed <u>M</u>etabolite <u>S</u>et <u>E</u>nrichment <u>A</u>nalysis (MSEA) (32) to evaluate the potential enrichment of functionally related metabolites in various metabolic pathways. The top MSEA-identified LCD-altered pathways at M3 included propanoate metabolism (Fig 3A), which includes several ketone body metabolites (i.e. 3-hydroxybutyric acid, 3-hydroxy-2-methylbutyriuc acid) noted to be increased by LCD (Fig 2). The second enriched pathway was porphyrin pathway that involves the metabolisms of porphyrins, such as heme or bilirubin (Fig 3A). In addition, LCD affected several amino acid pathways, including increased branched-chain amino acid biosynthesis (valine, leucine and isoleucine) as well as affected glycine, serine and threonine metabolism. Other listed pathways included selenocompound metabolism and aminoacyl-tRNA biosynthesis (Fig 3A). The top altered pathways at M6 in the LCD arm included phenylalanine, tyrosine and tryptophan biosynthesis, arginine and proline metabolisms and branched-chain amino acid biosynthesis (valine, leucine and isoleucine) (Fig 3B). The enrichment of these pathways are consistent with the changes of several amino acids, such as glycine, by the LCD at M6.

**Figure 3:**
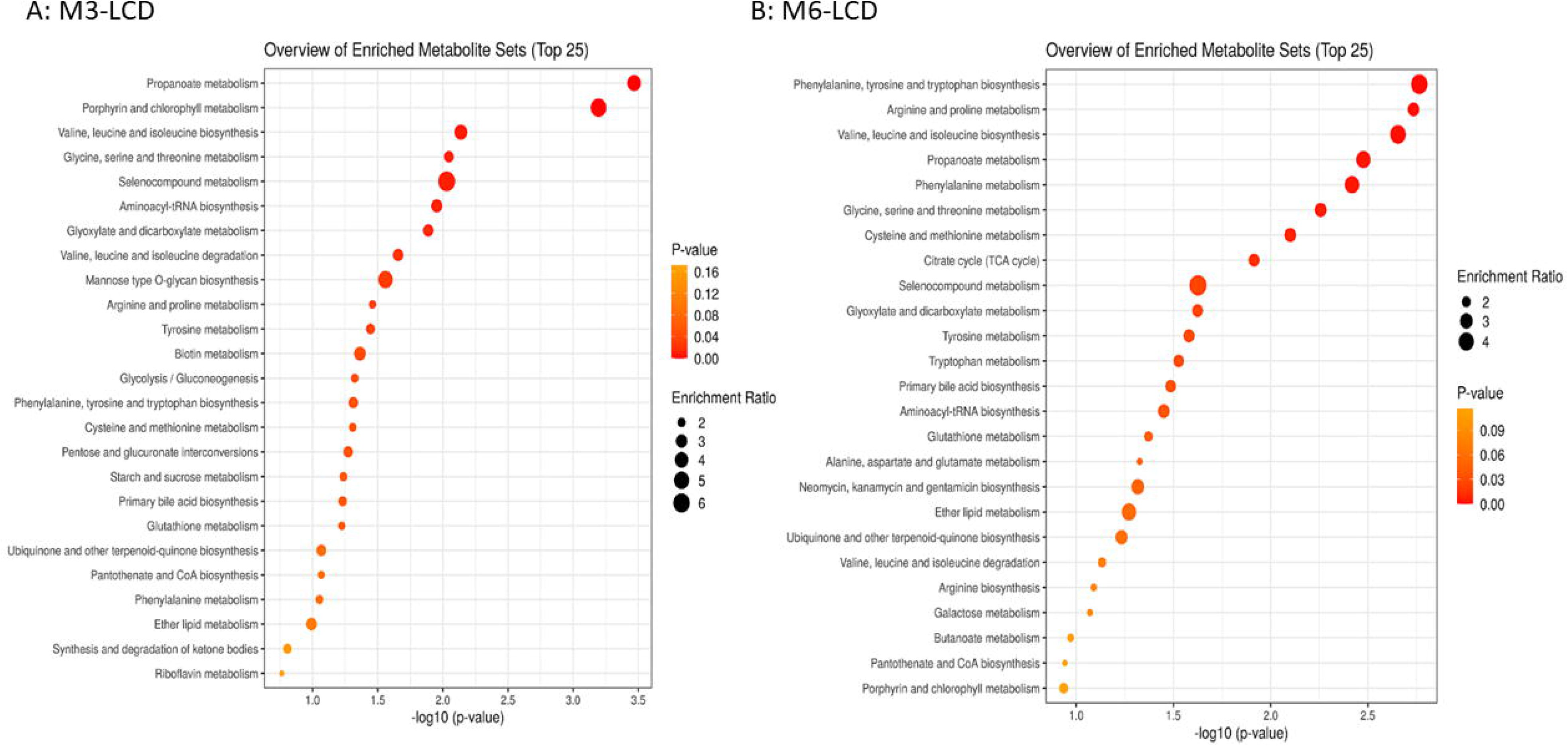
The enrichment of metabolic pathways by LCD at M3 and M6. Pathways Enrichment Analysis of the LCD-Affected metabolic pathways by MSEA at BL vs. LCD at M3 (A) and M6 (B).

### LCD-induced metabolic changes in both M3 and M6

To identify metabolites whose changes under LCD were statistically significant at both M3 and M6, we applied ANOVA - Simultaneous Component Analysis (ASCA module, MetaboAnlyst 4.0) to uncover the major patterns associated with the time points and the treatments. LCD significantly increased the level of several ketone metabolites, including 3-hydroxybutric acids (Fig 4A), hydroxybutyryl carnitine (Fig 4B) and 3-hydroxysuberoyl-carnintine (Fig 4C). These changes in ketone bodies were expected from LCD and indicate the adherence of the patients to the LCD intervention. In addition, LCD also increased the levels of glycine, and isobutyryl-acids (Fig 4D, E). Interestingly, there is also an increase in the hydroxy-isocaproic acid (HICA) at M3, a metabolite intermediate of leucine metabolism (Fig 4G). In addition, we calculated the changes of metabolites during LCD at M6 for each participant (Supplemental Fig 1A). These changes were then arranged by hierarchical clustering and the resultant heatmap shows the induction of all ketone bodies were co-clustered with glycine, 2-hydroxy-butyric acids, glycine and isobutyl-glycine, suggesting the coordinated increase of these metabolites under LCD (Supplemental Fig 1B).

**Figure 4:**
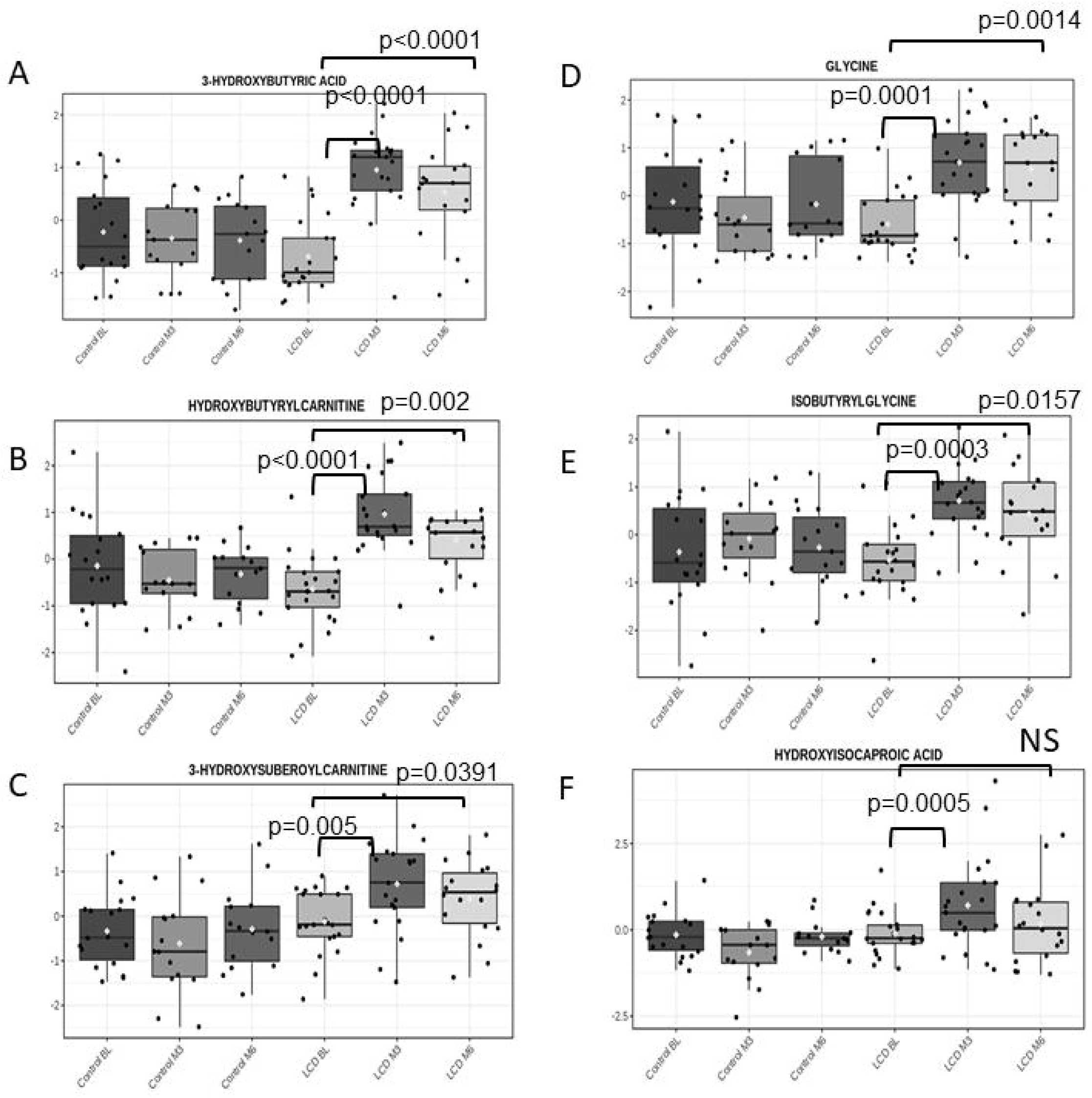
The metabolites which were significantly elevated by LCD. The effects of LCD-induced changes in the indicated metabolites at M3 and M6 in the control and LCD arms. The statistical significance (p values) of LCD-induced changes of indicated metabolites is shown.

In contrast, LCD also reduced the levels of alanine (Fig 5A), cytidine (Fig 5B), asymmetric dimethylarginine (ADMA) (Fig 5C). LCD also led to a significant reduction in the AICAR-cyclic-phosphate (Fig 5D), a cyclic nucleotide derivative of AICAR which is an exercise mimetic known to activate AMP-activated protein kinase (AMPK)(33). Finally, there is a significant reduction in the 2-oxobutanoate at M3 and even greater reduction at M6 (Fig 5E). Together, ANOVA identified many metabolites which were consistently altered by the LCD.

**Figure 5:**
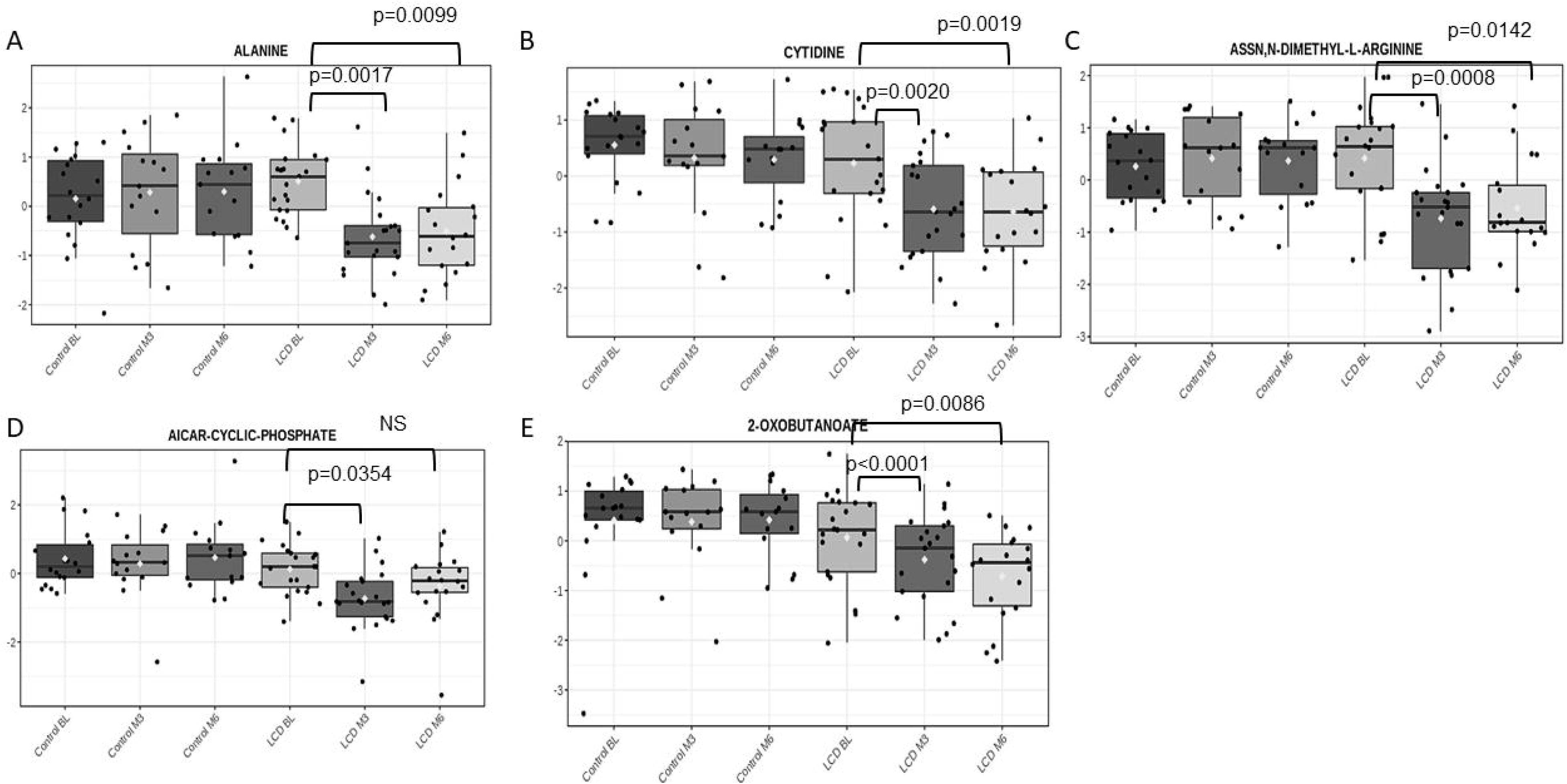
The metabolites which were significantly reduced by LCD. The effects of LCD-induced changes in the indicated metabolites at M3 and M6 in the control and LCD arms. The statistical significance (p values) of LCD-induced changes of indicated metabolites is shown.

### Serum metabolites whose LCD-affected changes correlate with PSA doubling time (PSADT)

In the primary paper describing the clinical outcomes of the CAPS2 trial, LCD was shown to increase PSADT in post hoc analyses (13). To understand the LCD-affected metabolic changes associated with longer PSADT, regression analysis was performed to calculate the correlation coefficients and associated p-values of each metabolite from Spearman correlation (non-parametric) to avoid the potential influence by outliers (Supplemental Table 3). Since LCD robustly increased the levels of ketone bodies, we compared the degree to which the increase in ketone bodies was associated with changes in PSADT. We focus specifically on the M6 because PSADT was measured using PSA values from the BL, M3, and M6 and thus we sought to assess metabolites whose change during the entirety of the study correlated with PSADT. Longer PSADT was significantly associated with higher 3-hydroxy-2-methylbutyriuc acid (Fig 6A), hydroxyl-butyrylcarnitine (Fig 6B), 2-hydroxybutyric acid (Fig 6C) at M6. There was also a strong trend between 3-hydroxy-butyric acid and PSADT (Fig 6D). Such positive correlation between the increase in the ketone bodies and longer PSADT raises the hypothesis that a greater ketogenesis under LCD may be associated with a slower tumor growth. Other than ketone bodies, PSADT was also correlated positively with the LCD-affected malate (Fig 6E) and citrate (Fig 6F), both of which were metabolites in tricarboxylic acid (TCA) cycle. PSADT also correlated with S-2-hydroxyethyl-N-acetylcysteinly-carnitine (Fig 6G), an acylcarnitine derivative of N-Acetyl-S-(2-hydroxyethyl)-L-cysteine, a product of detoxification of several environmental toxins. Reciprocally, PSADT showed a significantly negative correlation with the LCD-affected fructose-1-6-biphosate, a glycolysis metabolite (Fig 7A) and nicotinamide (Fig 7B). In addition, there was a negative correlation of PSADT with the changes in 2-oxobutanoate (Fig 7C) and octanoic acid (Fig 7D). We previously showed that androgen sulfate was the metabolite most consistently affected by ADT (19, 20), which is well-established as the main driver of PC growth. However, changes of androgen sulfate were not associated with PSADT in the current analysis (supplemental Figure 2). Taken together, these results suggest that a slower tumor progression and longer PSADT in the LCD arm is associated with more pronounced increased ketogenesis, TCA metabolites as well as more reduction in the fructose-1-6-biphosate, nicotinamide, 2-oxobutanoate and octanoic acid.

**Figure 6:**
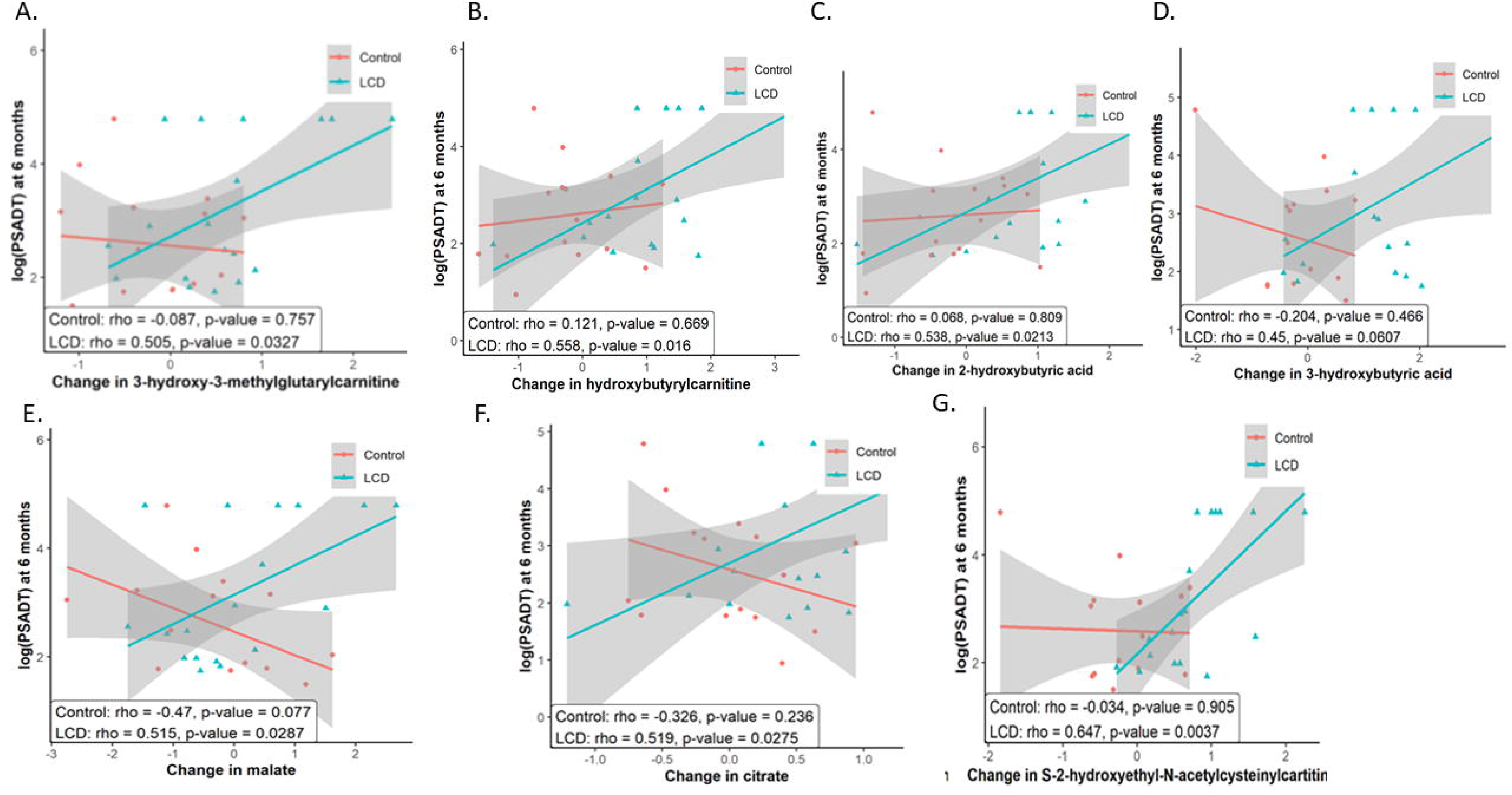
The metabolites whose LCD-changed positively correlated with PSADT. The correlation between the LCD-induced changes of indicated metabolites with PSADT in the control and LCD arms. The statistical significance (p values) of the correlation between indicated metabolites and PSADT is shown.

**Figure 7:**
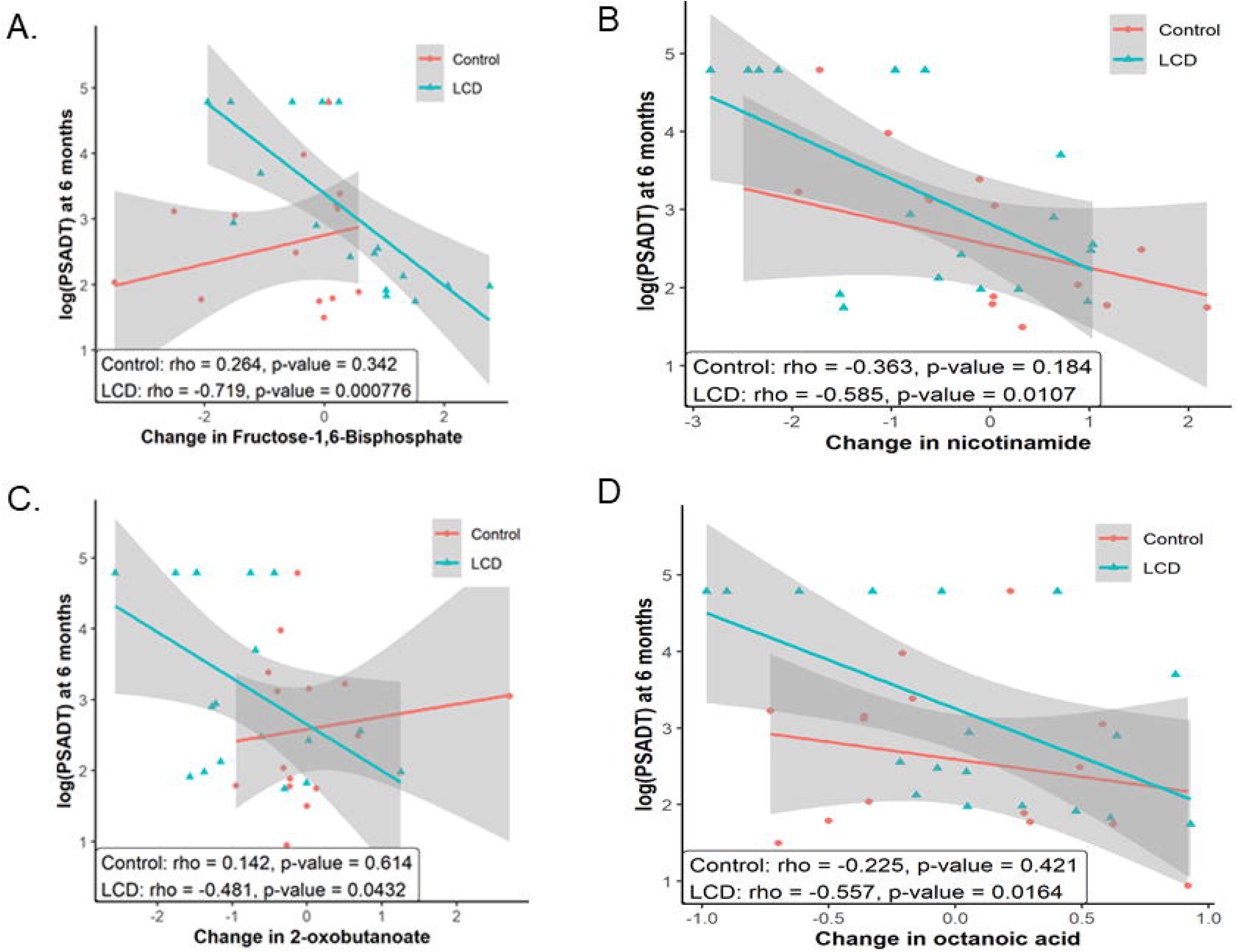
The metabolites whose LCD-changed negatively correlated with PSADT. The correlation between the LCD-induced changes of indicated metabolites with PSADT in the control and LCD arms. The statistical significance (p values) of the correlation between indicated metabolites and PSADT is shown.

## Discussion

In this study, we performed serum metabolome analysis of PC patients following a LCD intervention for 6 months. Such analysis revealed that LCD increased the levels of several ketone bodies, confirming the dietary adherence and the expected ketogenetic effects of LCD. More pronounced increase in the ketone bodies was generally associated with an increase in PSADT, implying a slower tumor progression. In spite of these general trends, only the increase of 3-hydroxy-2-methylbutyriuc acid. hydroxybutyrl-carnitine and 2-hydroxybutyric acids by LCD was significantly associated with PSADT. Small sample size or other confounding factors may have influenced the findings. Nonetheless, within the limits of a small study, and paralleling mouse data which found greater ketosis among mice on a ketogenic data were linked with better survival(11), these data raise the hypothesis that a ketogenic diet may delay PC growth. Ultimately, these intriguing results need to be further confirmed in future larger prospective randomized trials.

Other than ketone bodies, we also noted that LCD significantly reduced ADMA, an endogenous competitive inhibitor of nitric oxide (NO) synthase (34). Increased levels of ADMA are known to be associated with endothelial cell dysregulation, cardiovascular diseases and metabolic diseases (35, 36). Therefore, such LCD-reduced ADMA levels and expected benefit for endothelial cell function may contribute to the health benefit of LCD. Consistently, the levels of ADMA have been found to be affected by diets, including alcohol use and fruit and vegetable-rich diets (37, 38). Our results suggest that LCD may be another important means to reduce ADMA and improve endothelial cell function, though this requires further confirmation.

LCD did not consistently reduce all glycolysis-related metabolites assessed. However, shorter PSADT was associated with the increased glycolysis metabolite (fructose-1-6-biphosate) and reduced TCA metabolites (citrate and malate) in serum. Fructose 1,6-bisphosphate is a metabolic intermediate in the glycolytic pathway, which has been reported to enhance oncogenesis by activating Ras (39) and protect cancer cells from oxidative stress-induced cell death (40). The preferential use of glycolysis of tumor cells has been termed Warburg effects as one of the cornerstone of tumor metabolism and indeed is a key basis for applying LCDs in cancer (41). These changes in tumor metabolisms will help selected tumors with particular genetic compositions(42). Our findings are consistent with the hypothesis that more prominent reduced glycolysis and increased TCA metabolites under LCD is associated with slower PC progression.

LCD was also found to reduce cytidine, a pyrimidine nucleoside comprised of a cytosine bound to ribose. Cytidine can be converted to uridine by cytidine deaminase (CDA) and participate in the RNA synthesis. CDA can also metabolize many chemotherapeutic agents and its serum levels and activities can be highly predictive of toxicities and side effects of chemotherapies (43). While we cannot conclude whether the lower serum cytidine level is associated with an increased CDA activities, previous reports have suggested that LCD can mitigate the toxicities associated with chemotherapies (44), consistent with a possible connection between LCD and low cytidine via regulation of CDA levels. Indeed, other studies have shown that fasting, which shares parallels with a LCD in terms of both being ketogenic, can reduce chemotherapy toxicity (33, 45). Whether an LCD has similar properties requires formal testing in the future.

We also observed significant changes in several metabolites whose biological connection to the relationship between LCD and PC progression is less clear. For example, LCD significantly reduced the levels of 2-oxobutanoate and the degrees of its reduction was associated with longer PSADT. 2-oxobutanoate, also known as 2-ketobutyric acid is a product of the lysis of cystathionine (46). Cystathionine, together with hydrogen sulfide (H2S), is formed by cystathionine β-synthase (CBS) during the trans-sulfation pathway. CBS has been found to promote the growth of many tumor types, possibly due to prevent ferroptosis (47), the signaling function of H2S (48, 49) and pro-growth effects of cystathionine (48). It is possible that the reduced levels of 2-oxobutanoate under LCD may reflect the increased turnover of cystathionine as well as the dysregulation of other amino acids, such as glycine and alanine. However, the significance and potential mechanism are unknown.

Our results should be interpreted within several limitations. First, a small sample size may limit our power to detect biologically relevant differences that may be of importance. As such, our results should be considered hypothesis-generating and require validation in larger datasets. We did not have complete metabolic data on all patients which may have disrupted our randomization balance. Furthermore, the small datasets make it difficult to analyze any potential differences for patients with different clinical characteristics (i.e. different races, tumor grades, etc.). In addition, the relative contribution of LCD itself and the weight loss outcome to the observed effects cannot be distinguished in this study and needs to be further dissected. Finally, future studies are needed to correlate these metabolomic changes with the long-term outcomes including tumor control and metabolic side effects such as diabetes and cardiovascular diseases.

## Funding and Acknowledgement

American Urological Association Foundation, BERG, NIH, and Robert C. and Veronica Atkins Foundation

## Author contributions statement

JTC: data analysis, manuscript writing – original draft, review and editing. PHL: Conceptualization, data analysis, manuscript writing. VT, VB, NRN, BG, RS: methodology, data acquisition, statistical analysis. TO: statistical analysis. EC: methodology, project, administration, data acquisition, statistical analysis. CGA and AR: manuscript writing. MAK: Conceptualization, methodology, project administration, data acquisition, statistical analysis. SJF: Conceptualization, funding acquisitions, manuscript writing.

## Data Availability Statement

The metabolomic data will be made available to the academic community upon publication of the manuscript.

---

**Supplemental Figure 1: Heatmap of the LCD-induced changes in all metabolites at M6** (A) The changes of all metabolites from BL in response to control and LCD arms at M6 were derived by zero-transformation and arranged by hierarchical clustering. Yellow indicates an increase, blue indicates a reduction and black indicates no change. (B) Clusters of metabolites that were induced by LCD in most samples were expanded with the names of the metabolites shown.

**Supplemental Figure 2: The correlative relationship between LCD-changed androsterone sulfate with PSADT at M6**

**Supplemental Table 1**: Top metabolites affected by the LCD at M3

**Supplemental Table 2**: Top metabolites affected by the LCD at M6

**Supplemental Table 3**: The serum metabolites ranked by the correlation with the LCD-induced changes with PSADT at indicated times of control and LCD arms.

